# Non-annual seasonality of influenza-like illness in a tropical urban setting

**DOI:** 10.1101/100222

**Authors:** Ha Minh Lam, Amy Wesolowski, Ngyuyen Thanh Hung, Tran Dang Nguyen, Nguyen Thi Duy Nhat, Stacy Todd, Dao Nguyen Vinh, Nguyen Ha Thao Vy, Tran Thi Nhu Thao, Nguyen Thi Le Thanh, Phan Tri Tin, Ngo Ngoc Quang Minh, Juliet E Bryant, Caroline O Buckee, Tran Van Ngoc, Nguyen Van Vinh Chau, Guy E Thwaites, Jeremy Farrar, Dong Thi Hoai Tam, Ha Vinh, Maciej F Boni

**Author notes:** Contributed equally (Ha Minh Lam, Amy Wesolowski, Ngyuyen Thanh Hung). Correspondence Maciej F Boni, Center for Infectious Disease Dynamics, Department of Biology, Pennsylvania State University, University Park, PA, 16802.

## Abstract

In temperate countries, influenza and other viral respiratory diseases often have distinct seasonal peaks occurring during colder, wintertime months. However, little is known about the dynamics of influenza and viral respiratory disease dynamics in the tropics, despite high morbidity and a clear epidemiological link between tropical and temperate countries. In temperate countries, the dynamics of influenza and other respiratory diseases are often analyzed using syndromic surveillance data describing influenza-like illness (ILI) as ILI is highly correlated with virological surveillance for influenza. To obtain a detailed picture of respiratory disease incidence patterns in a large tropical city, we established an mHealth study in community outpatient clinics in Ho Chi Minh City, Vietnam (11N latitude). From August 2009 through December 2015, clinics reported daily case numbers of ILI using standard mobile-phone SMS messaging. A subset of these clinics performed molecular diagnostics for influenza A and B viruses. Unlike the annual patterns seen in temperate countries, ILI activity in Ho Chi Minh City exhibited strong non-annual periodicity and was not correlated with PCR-confirmed influenza. The dominant periodicity in the data was approximately 200 days. This was confirmed by a time series decomposition, a step-wise regression analysis on annual and non-annual covariates, and a forecasting exercise showing that forecasting was 30% to 40% more accurate when a 200-day non-annual cycle was included in the forecast. This suggests, for the first-time, that a non-annual cycle may be an essential driver of ILI dynamics in the tropics. This raises new questions about the seasonality and drivers of respiratory disease transmission in tropical countries.

## 1 Introduction

In temperate countries, influenza virus is one of the most studied disease systems, exhibiting a predictable wintertime transmission season and a robust relationship between syndromic and molecular surveillance. Little is known about the epidemiology of influenza virus in the sub-tropics and tropics despite a renewed research interest in tropical influenza over the past decade resulting from increased availability of influenza surveillance and sequence data^1–5^. To date, research on tropical influenza has concentrated on whether influenza epidemics exhibit annual seasonality^6–14^ and whether influenza viruses show patterns of year-round persistence^15–19^. A third question that has received less attention is whether syndromic influenza-like illness (ILI) surveillance has the same peaks and troughs as molecular surveillance for influenza virus in these regions. In temperate countries, public health agencies are able to rely on ILI reporting to signal the onset of the influenza season^20–22^, but it is not known if ILI and influenza correlate in tropical countries^23,24^.

The majority of epidemiological studies looking at influenza and/or respiratory disease in the tropics have two major drawbacks. The first one is ignoring absolute case counts and reporting only the percentage of samples (nose/throat swabs) that test positive for influenza^12,14,24–27^. Ignoring case counts makes it impossible to determine if samples are being taken during a potential influenza season or outside of it. The second one is underpowering the analysis by using a short time series or monthly data or both^23,24,26–32^. Monthly data are normally too coarse to infer the presence of an annual transmission season or other periodic trends (if these exist) unless the time series is quite long. In fact, this is one of the reasons for disagreement in the current literature as some studies on respiratory disease in the tropics claim support for an annual transmission season^7,12,14,26–28,33–35^ while others show mixed or no evidence^8,13,31,36–40^. Among these, some of the more weakly supported results are being used in public health policy to advocate for particular vaccination timings based on incorrectly identified seasonal signals^12,35^.

Understanding the dynamics of tropical influenza – especially the presence or absence of seasonality – may allow the forecasting methods successfully deployed in temperate countries^41,42^ to be used for tropical influenza. Current forecasting methods rely on mechanistic Susceptible-Infected-Recovered (SIR) models and known/inferred climate associations to accurately predict increases in influenza virus infections. In the tropics, it is not known whether influenza dynamics obey classic SIR models, whether they are characterized by low-level persistence, or a combination of the two. It is also not known which climate-influenza associations are expected to be present in tropical countries despite some recent advances on this topic^11,43^. If the intrinsic epidemiological dynamics and the presence/absence of climate associations can be understood, forecasting of influenza epidemics in the tropics may be possible. Thus far, the only attempt at influenza forecasting for the tropics or subtropics reported that the majority of forecast attempts (different leads, different methods) had accuracies below 50% when predicting the timing of an influenza peak^44^.

In addition, an accurate description of the basic epidemiology of tropical influenza is critical for inferring the likely routes of viral seeding from the tropics to temperate zones and vice versa^4,45^. Although there is abundant phylogeographic evidence linking tropical and temperate influenza sequences^1^, very few analyses have investigated the epidemiological characteristics of tropical influenza and how these affect epidemics in temperate zones. Two exceptions can be seen in Brazil and China, both of which span multiple climatic zones. In Brazil, a pneumonia and influenza mortality time series dating back to 1979 shows an annual influenza epidemic progressing from tropical to temperate parts of Brazil^17^. A second example can be seen in a recent study published using sentinel surveillance data from in China, showing the transition from large wintertime influenza peaks in the north to smaller less predictable peaks in the subtropics^46^. Beyond these two examples, epidemiological links between the tropics and other regions are hard to show due to the paucity of long-term consistent surveillance data in tropical regions.

To investigate the fine-scale epidemiology of respiratory disease dynamics in the tropics and evaluate the potential for forecasting, in August 2009 we set up a real-time community-based participatory epidemiology network in Ho Chi Minh City, Vietnam. Our initial hypothesis was that influenza-like illness trends in Ho Chi Minh City would not be annual. Enrolled outpatient clinics across the city reported daily case numbers of influenza-like illness by standard mobile-phone SMS messages. A subset of the clinics provided molecular confirmations of influenza virus in order to assess the relationship between ILI and influenza. Our goals were to make daily reporting of influenza-like illness as simple as possible in order to encourage frequent reporting and wide participation, and to create a real-time ILI surveillance system that could be used by health professionals in Ho Chi Minh City. Our study is most similar to the clinic-centered mHealth systems set up in Senegal^32^ and Madagascar^47^, and the benefits of this type of real-time, big-data epidemiology can be seen in the dengue hotline system recently described by Rehman et al^48^. The purpose of our study was to build a long-term consistent time series of both ILI reports and influenza molecular confirmations. We analyzed the data with traditional time series decomposition to detect periodic signals, with stepwise regression analyses to determine the importance of climate and other covariates, and with regression-based forecasting to determine the predictability of ILI trends in Ho Chi Minh City.

## 2 Results

A total of 63 clinics were enrolled during the study, about half of which reported regularly, and 37,676 daily reports were received from August 10, 2009 to December 31, 2015, corresponding to 1,759,473 outpatients and 186,346 outpatients meeting the clinical definition of influenza-like illness. The median clinic saw an average of 30 patients per day (IQR: 16–50 across clinics). Approximately 10.6% of all patients were classified as ILI, and this percentage exhibited a decreasing trend during the first six years of the study (Table 1). To create a single ILI time series for Ho Chi Minh City, we detrended and standardized each clinic’s ILI percentages to a z-score scale and then aggregated these into a single z-score time series. Several internal validations were done to ensure that the data followed certain expected behaviors for multi-site syndromic reporting and that arbitrary or random reports were not being sent during the course of the study (see Materials and Methods). In particular, note that individual clinic time series correlated with each other, and replacing a single clinic with a white noise signal of equal variance reduced the correlation between that clinic and the aggregate ILI trend (Figure S2). Influenza-like illness trends in Ho Chi Minh City (Figure 1) suggest that there are typically multiple ILI peaks per year, as has been observed in other tropical and sub-tropical regions^11,17,44^. Visually, no seasonal or annual cycle appears in these data.

**Table 1:** Summary of ILI reports for 2009-2015

| Year | Clinics reporting at least |  |  | Total Patients | Reported ILI Cases | ILI Percentage |  |
| --- | --- | --- | --- | --- | --- | --- | --- |
|  | 1 day | 50 days | 150 days |  |  | Median | IQR |
| 2009* | 19 | 10 | 0 | 35,115 | 10,163 | 24.40 | 19.36 , 35.89 |
| 2010 | 27 | 15 | 7 | 103,396 | 24,922 | 15.42 | 3.82 , 26.89 |
| 2011 | 28 | 24 | 20 | 275,033 | 35,176 | 14.73 | 4.13 , 25.86 |
| 2012 | 35 | 28 | 25 | 375,077 | 42,373 | 13.30 | 6.25 , 26.49 |
| 2013 | 30 | 28 | 23 | 385,300 | 30,183 | 9.99 | 2.47 , 20.61 |
| 2014 | 32 | 27 | 20 | 300,223 | 19,461 | 10.64 | 2.67 , 16.14 |
| 2015 | 35 | 26 | 21 | 252,932 | 21,318 | 11.69 | 6.86 , 17.58 |

\* Data collection in 2009 started on August 10th.

**Figure 1.**
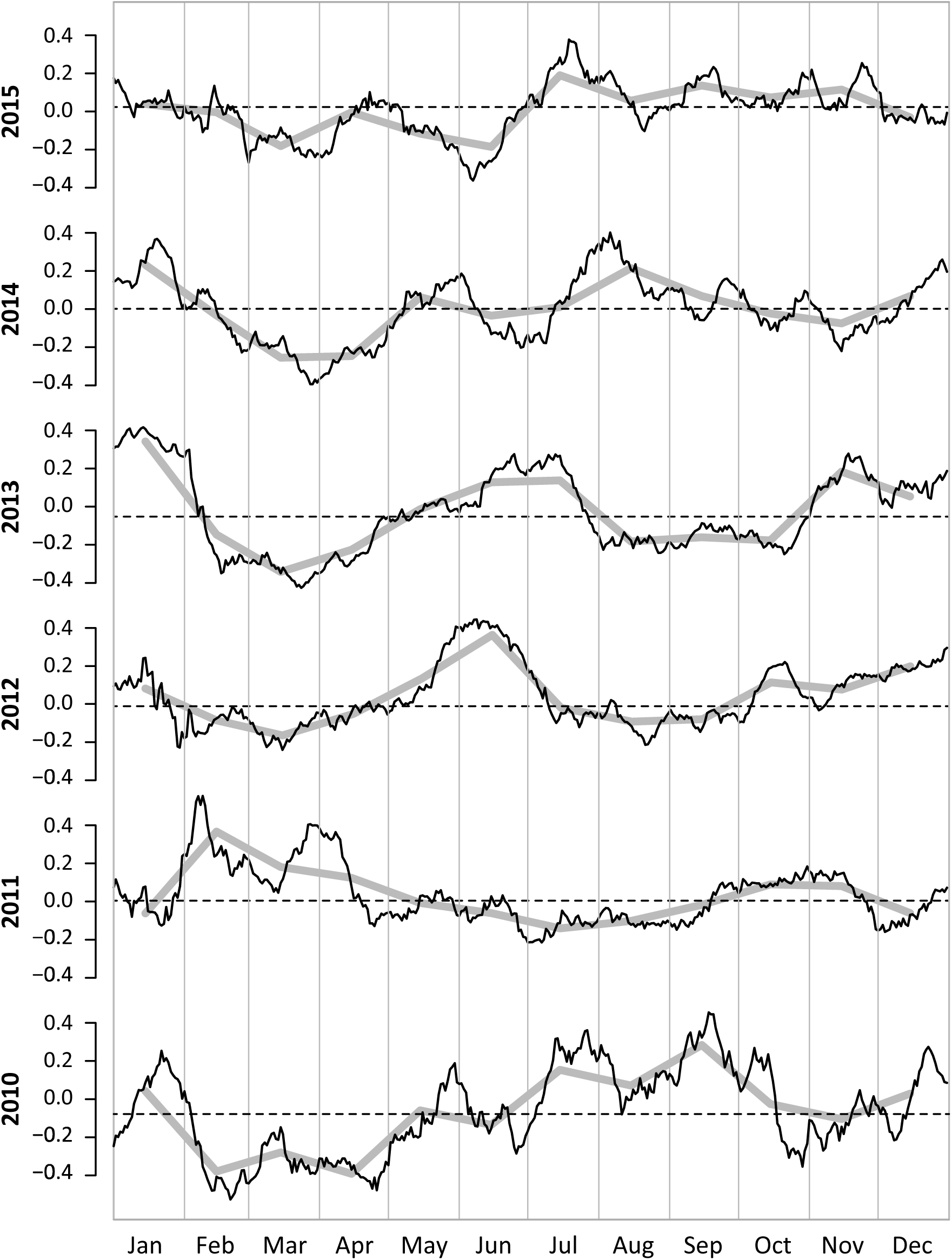
Trends in ILI z-scores by year. The black lines show 15-day moving-average smoothed z-scores (after detrending). The gray solid lines show the monthly mean z-score values. The horizontal dashed lines represent the median ILI z-score for that year.

In a subset of the clinics, molecular confirmations on naso-pharyngeal samples (*n* = 2,217) were taken from May 2012 to December 2015. Compared to other tropical settings, these clinics had a rate of influenza positivity (21.5% positivity for influenza A and 9.7% positivity for influenza B) in the high range of previously published studies^14,23,28,36,37,49^. We compared the confirmed influenza cases to the ILI data and found that there was no correlation between the two time series (Figure 2; Pearson correlation coefficient: −0.02, p-value: 0.86) and that this did not differ for influenza A and B individually (both p-values > 0.15). The time series showed periods of high ILI activity with a low level of influenza confirmation, likely representing epidemic waves of other respiratory viruses, as well as periods that were high influenza and low ILI, suggesting that an influenza epidemic may not contribute as much to the overall ILI trend as it does in temperate regions.

**Figure 2.**
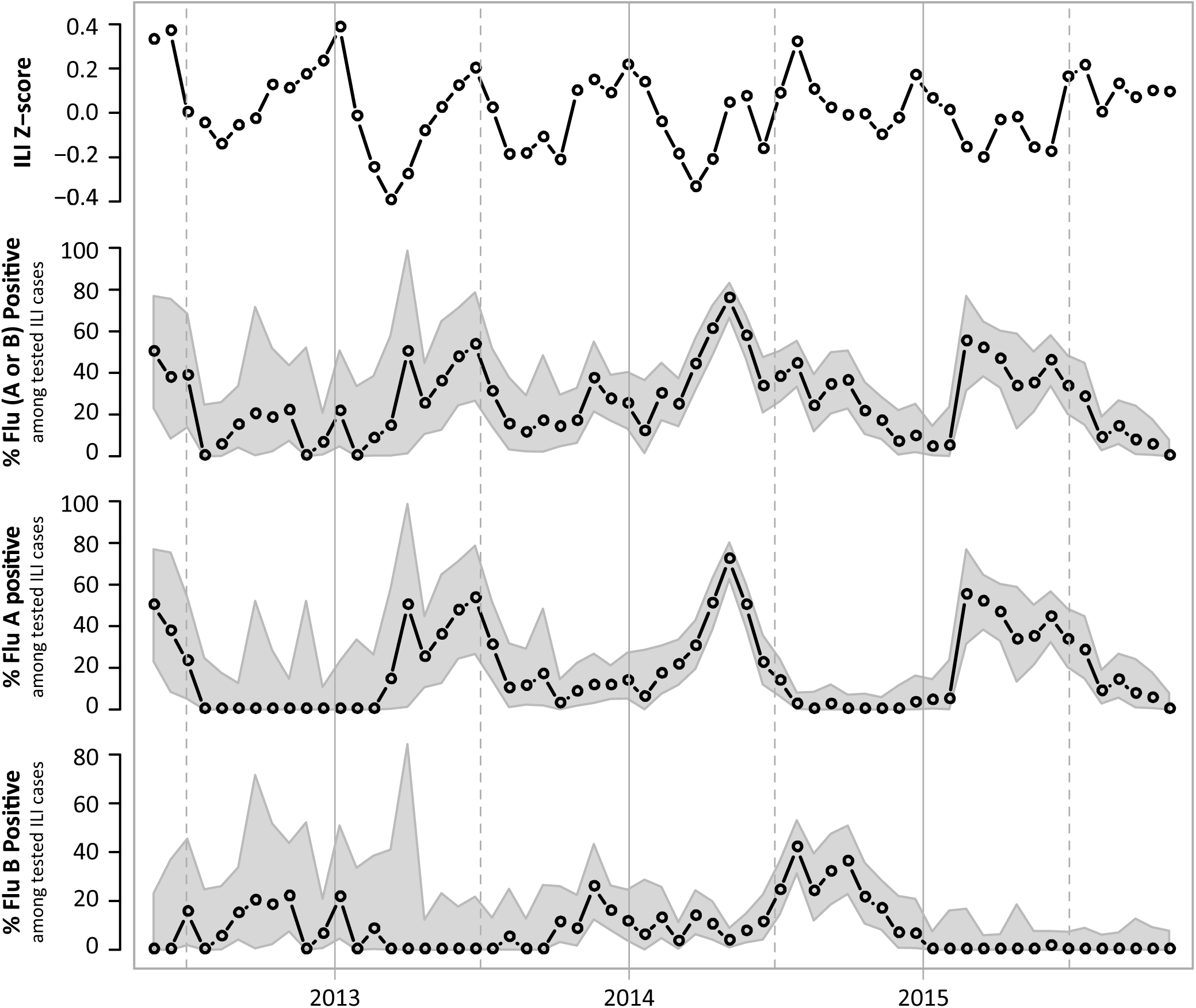
Time series of ILI z-score and influenza PCR-positivity, in 3-week windows, for the period of time when PCR confirmations were being done in the clinics in the study. Gray region around flu-positive percentage is the 95% confidence region computed using the exact binomial method. The Pearson’s correlation between the time series is shown in Table S2.

We identified a dominant periodicity in the data using an auto-correlation function and standard time series decomposition (see Materials and Methods). The auto-correlation function (ACF) identified 206 days (*ACF* = 0.262; p-value < 10^−15^) whereas the discrete Fourier transform identified 199 days as the time series’ dominant periodic signal (*ACF* = 0.244 for a lag of 199 days; p-value < 10^−15^); see Figure 3. This non-annual signal is almost twice as strong as the annual cycle, with the 365-day lag exhibiting an auto-correlation value of 0.153 (p-value = 0.014); note that the large number of data points results in statistical significance for nearly all ACF values. A dominant non-annual signal is an unusual feature in disease incidence data. We verified that this result was not an artifact of our data renormalization and detrending methods by applying these same methods to temperate zone ILI data and showing that ILI time series in Europe and North America show their strongest periodic signals at 365 days, with no evidence of periodic signals shorter than one year (Figure S3).

**Figure 3.**
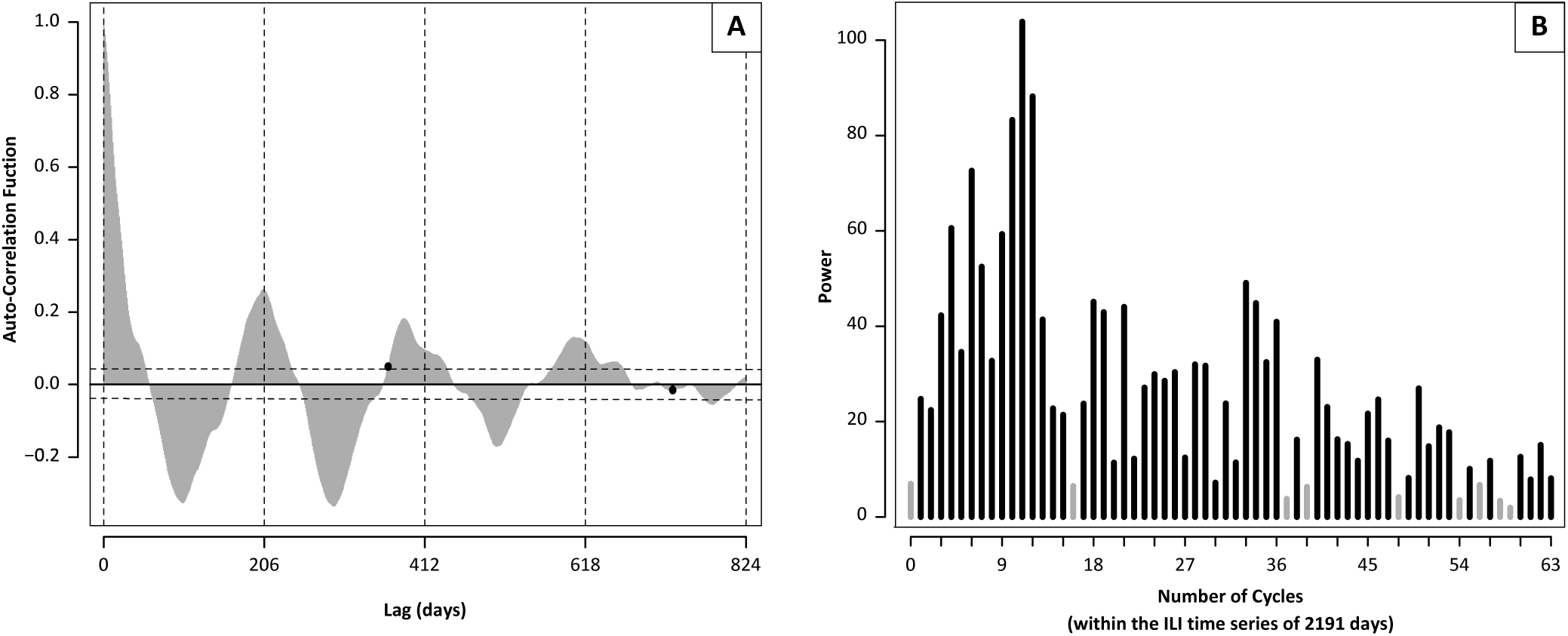
(**A**) Autocorrelation function (ACF) for the z-score time series. Horizontal dashed lines demark the statistically significant regions (*p* < 0.05). Black dots represent the ACF values of lags of 365 and 730 days. The first peak in the ACF is at the lag of 206 days. (**B**) Discrete Fourier transform (DFT) of the z-score time series. The period length of each DFT can be calculated by dividing 2,191 (the number of days in the time series) by the corresponding number of cycles (the frequency of the DFT). Frequencies whose power is lower than 6.93 (i.e. periodic functions whose correlation with the z-score time series is lower than their correlation with a constant signal) are shown in gray. The DFT reaches its highest power at 11 cycles, corresponding to a cycle length of 199 days.

To determine the relative influence of annual and non-annual signals on the ILI trend, we performed a stepwise regression of the ILI trend onto both annual climatic variables and the system’s intrinsic non-annual cycle. Lagged variables, interactions, and non-linear transformations of the climate variables were included; the non-annual cycle was constructed as a step-function with periodicity 206 days (see Materials and Methods). The stepwise regression indicated that the terms with explanatory power were the daily temperature, relative humidity (RH and √RH), the interaction term between RH and temperature, lagged climate terms (see eq. 2 in Materials and Methods), and the non-annual cycle. Clearly, the association between the non-annual effect and the ILI trend is statistically significant (Table 2), and the non-annual effect is identified using Akaike Information Criterion as a component of the best fit model. Nevertheless, it is important to remember that the number of data points (>37,000) results in statistical significance for a large number of annual and non-annual covariates. Thus, additional robustness analyses were performed.

**Table 2:** Estimates of coefficients from regressing the smoothed daily ILI z-scores (2010-2012) onto two climate variables, an interaction term, and the temporal indicator variables that were used to construct a periodic 206-day forcing function in the time series. Temperature was measured in Celsius.

|  | Coefficient | Estimate | Std. Error | t statistic | P-value |
| --- | --- | --- | --- | --- | --- |
| Intercept |  | 63.5198 | 6.1223 | 10.3751 | 4.31E-24 |
| Temperature |  | -1.0446 | 0.0719 | -14.5345 | 8.43E-44 |
| 3-week Lagged Temperature |  | 0.0004 | 0.0128 | 0.0338 | 9.73E-01 |
| 4-week Lagged Temperature |  | -0.0232 | 0.0180 | -1.2894 | 1.98E-01 |
| 5-week Lagged Temperature |  | 0.0843 | 0.0120 | 7.0508 | 3.19E-12 |
| Rel. Humid. |  | -0.0584 | 0.0375 | -1.5569 | 1.20E-01 |
| Sq. Root Rel. Humid. |  | -5.3047 | 0.7488 | -7.0839 | 2.54E-12 |
| 5-week Lagged Rel. Humid. (RH.lag5) |  | 0.1656 | 0.0412 | 4.0184 | 6.27E-05 |
| Sq. Root RH.lag5 |  | -2.8670 | 0.7278 | -3.9393 | 8.70E-05 |
| Rel. Humid. × Temperature |  | 0.0128 | 0.0009 | 13.5127 | 1.60E-38 |
| Temporal Interval |  |  |  |  |  |
| 2 |  | 0.0098 | 0.0275 | 0.3576 | 7.21E-01 |
| 3 |  | -0.0963 | 0.0274 | -3.5128 | 4.62E-04 |
| 4 |  | -0.1900 | 0.0275 | -6.9097 | 8.34E-12 |
| 5 |  | -0.2111 | 0.0275 | -7.6857 | 3.44E-14 |
| 6 |  | -0.2178 | 0.0282 | -7.7159 | 2.75E-14 |
| 7 |  | -0.1105 | 0.0274 | -4.0262 | 6.07E-05 |
| 8 |  | -0.1036 | 0.0279 | -3.7205 | 2.09E-04 |
| 9 |  | -0.1442 | 0.0275 | -5.2471 | 1.86E-07 |
| 10 |  | -0.0867 | 0.0278 | -3.1233 | 1.84E-03 |
| 11 |  | -0.1811 | 0.0293 | -6.1726 | 9.53E-10 |
| 12 |  | -0.1949 | 0.0280 | -6.9524 | 6.25E-12 |
| 13 |  | -0.2311 | 0.0279 | -8.2917 | 3.33E-16 |
| 14 |  | -0.0828 | 0.0274 | -3.0191 | 2.60E-03 |
| 15 |  | 0.0290 | 0.0275 | 1.0569 | 2.91E-01 |
| 16 |  | -0.0563 | 0.0267 | -2.1069 | 3.54E-02 |
| 17 |  | 0.0047 | 0.0262 | 0.1790 | 8.58E-01 |
| 18 |  | 0.0347 | 0.0265 | 1.3107 | 1.90E-01 |
| 19 |  | 0.0072 | 0.0263 | 0.2750 | 7.83E-01 |
| 20 |  | -0.0101 | 0.0260 | -0.3873 | 6.99E-01 |
| 21 |  | 0.0548 | 0.0268 | 2.0440 | 4.12E-02 |

As a third validation of the existence of a non-annual cycle as a true feature of respiratory disease transmission in Ho Chi Minh City, we tested the sensitivity of the ILI forecast accuracy to the length of the non-annual cycle and to the amplitude of the trends of climate variables. The rationale is that if an intrinsic non-annual cycle truly influences respiratory disease dynamics, then (1) forecasting of respiratory disease should be possible using the non-annual cycle, and (2) the forecasts should be less accurate if the non-annual cycle is not used or if an artificial non-annual cycle of a different periodicity is used. Regressing the 2010-2012 portion of the time series onto the AIC-selected covariates (including the non-annual cycle of length *c* = 206), we were able to predict the 2013-2015 ILI time series with a median absolute error of 0.129 on a z-score scale (Figure S6A). A sensitivity analysis indicated that forecast accuracy is very sensitive to the intrinsic cycle length, and it is reduced substantially if the length *c* of the non-annual cycle is changed by a small amount (Figure 4); the median prediction error is approximately 40% to 50% higher when forecasting is performed with a cycle length *c* < 195 or *c* > 215. The increase in prediction error is small or non-existent when the climate variables are smoothed to reduce their correspondence with the true climate time series (Figure 4). Thus, the non-annual cycle is the key characteristic of this dynamical system that enables accurate forecasting.

**Figure 4.**
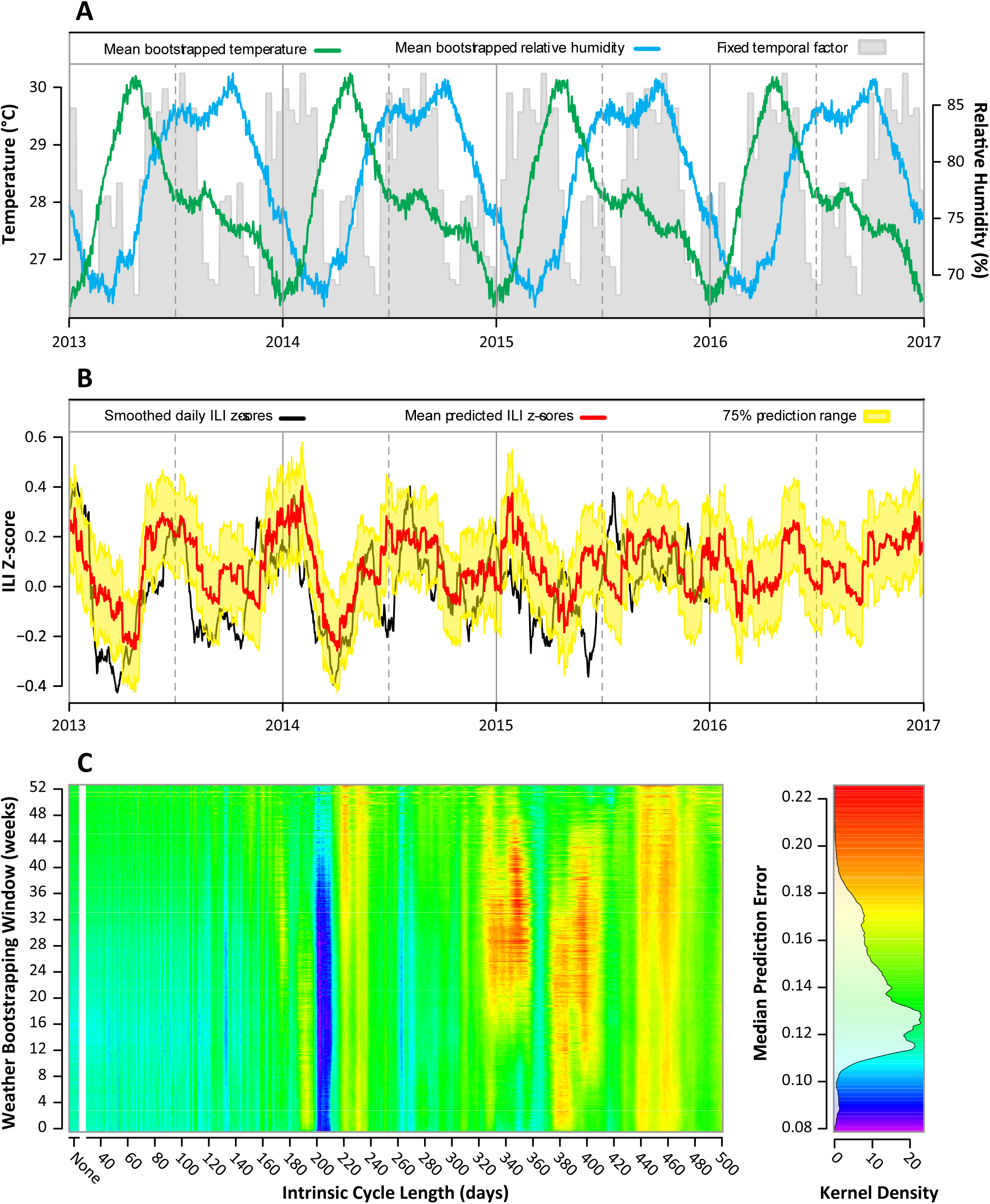
Forecasting ILI z-scores with bootstrapped weather data. (**A**) Annual average temperature trend (green) and relative humidity trend (blue) based on 2000-2015 weather data for Ho Chi Minh City. Bootstrapping is done in a 21-day window around each time point, which has the effect of smoothing the data with a 21-day window. The shaded gray area shows the inferred periodic signal from equation (2) using the 2010 to 2012 z-scores and assuming a 206-day cycle. (**B**) Predicted daily ILI z-scores from the regression model (red) and their 75% prediction range (yellow) are plotted alongside with the daily ILI z-scores (black). Model parameters were estimated by regressing ILI z-scores of 2010-2012 on the real weather data of 2010-2012. Predictions were calculated based on bootstrapped weather data (see Materials and Methods). The median prediction error from January 1 2013 to December 31 2015 is 0.125 (z-score scale, IQR: 0.064, 0.203). (**C**) Median prediction errors when varying both the width of the bootstrapping window *d* for the weather data and the duration of the intrinsic cycle *c* in the system (see Methods). The minimum prediction error is achieved with a weather bootstrapping window of 199 days and an intrinsic cycle of 202 days.

Several robustness tests were performed. Figure S6 shows that forecasting using a 202-day intrinsic non-annual cycle in combination with bootstrapped climate data gives the most accurate forecasts, and that a 211-day cycle was optimal when forecasting ILI trends using real weather data. These results are robust to whether mean or median prediction error is used as an evaluation criterion (Figure S7). Using a simpler regression model with no lags and no non-linear climate terms, a 201-day cycle gave the lowest prediction errors (Figures S8 and S9). All analyses provided support for the existence of a non-annual cycle with periodicity of approximately 200 days.

Our decomposition, stepwise regression, and prediction analyses provide strong evidence that an intrinsic non-annual cycle of around 200 days exists for respiratory disease transmission in Ho Chi Minh City. This cycle is either unique to the dynamics of respiratory infections in tropical climates, or it is a natural part of respiratory disease epidemiology in all regions but not detectable in temperate countries as a result of being overwhelmed by the strong winter seasonality of respiratory disease transmission. An ILI indicator, showing whether ILI percentages are above or below the mean trend, is updated daily and publically available (www.ili.vn) providing a real-time surveillance system for patients and clinical providers.

## 3 Discussion

Our study demonstrates the value of community epidemiology studies for describing fine-scale dynamics of influenza-like illness in tropical settings where respiratory disease dynamics are non-annual and difficult to predict. We were able to show that a network of community clinics can generate a high-quality syndromic time series that can be used to understand local transmission patterns of respiratory disease, and that such a network can generate a significantly larger data set (~6,000 data points per year) than traditional surveillance systems that report weekly or monthly measures of incidence. This volume of data increases statistical power to detect ILI associations, and in our study, the presence of non-annual forcing in the system. The present study does not achieve the data volume seen in ‘big data’ study designs^21,50–52^ which can have tens of millions of observations per year, but the specificity of our data signal is higher than in the aforementioned studies as each data point in our study corresponds to a patient, seen by a physician, determined to have met or not met the clinical criteria for influenza-like illness.

The major quality control challenge we encountered was accounting for long-term trends in ILI (we had a downward trend in our data). In a multi-site time series, detrending must be done carefully, and changes in a site’s reporting patterns must be investigated individually. From discussions with the reporting physicians in our study, the putative causes of the decreasing trend in ILI were likely to have been (*i*) a more than doubling of patient visit costs that would have reduced the likelihood of reporting a minor respiratory illness, (*ii*) increased clinical specialization at some sites, or (*iii*) more conservative interpretation of ILI guidelines after molecular diagnostics were introduced in May 2012. In addition, during 2011 and 2012 a few large clinics were enrolled in the study, and some of these had higher patient volumes but lower ILI percentages. All of these features of community-based syndromic reporting systems need to be considered for both study design and surveillance purposes. Detrending with a 12-month moving average appears to be the simplest way to detrend and preserve any potential annual structure in the data.

The lack of correlation between influenza trends and ILI trends suggests that the transmission dynamics of respiratory disease differ between tropical and temperate zones, consistent with the past decade’s literature on this topic^10,11,13,17,43,46^. Given the observed pattern of multiple ILI peaks in our data, some of which are influenza epidemics and some of which are not, the natural hypothesis explaining this pattern is that multiple respiratory pathogens co-circulate and cause asynchronous epidemics. It is unknown if in such a system multiple respiratory pathogens should circulate independently or not. The putative mechanism that would create dependence or interference among waves of different co-circulating respiratory viruses would be post-infection raised antibody or cytokine concentrations^53–55^ generated by one viral epidemic preventing an epidemic of a different virus from taking off immediately thereafter. Epidemiological interference among respiratory viruses has been observed in long-term time series in temperate^56,57^ and tropical^58^ regions, but the strength and duration of this effect is not well understood. In our community study, additional molecular confirmations for a range of respiratory pathogens are now underway to further describe this phenomenon.

The second major question that arises from the basic correlational analysis between ILI and influenza is why high influenza periods should be observed when ILI is low. To the best of our knowledge, this pattern has not been observed in other surveillance systems, as a wave of influenza infections is normally sufficient to generate a substantial uptick in the ILI signal. The likely explanation for a high-influenza low-ILI period is a larger than expected prevalence of other respiratory viruses among the reported ILI cases; this is possible as the community clinics in our study are almost exclusively outpatient and likely to see many mild cases of respiratory disease. If influenza infection represents only a small fraction of respiratory disease among these outpatients, a wave of influenza alone would not generate an ILI peak. In general, community-based studies of respiratory disease should aim to characterize the contribution of all respiratory viruses to the ILI trend to determine if it is a particular pathogen’s dominance or synchrony among certain pathogens that generates an ILI peak.

The major finding in our study is that the dominant periodicity observed in our ILI time series is non-annual. This is the first report of a non-annual disease cycle in temperate or tropical respiratory disease data. The existence of an intrinsic non-annual cycle in the dynamics is supported by traditional time series decomposition, by a regression of the time series onto both annual and non-annual covariates, and by an analysis of the system’s predictability showing that accurate forecasts of ILI trends are highly dependent on the system’s non-annual cycle of ~200 days. The presence of non-annual periodicity is consistent with a mechanism of short-term non-specific immunity conferred by one respiratory virus that affords near-term protection (3-6 months) against infection with other respiratory viruses. Data on the rate of antibody decay after acute influenza infection are consistent with this hypothesis^54,55^, but unfortunately no such data exist for other respiratory viruses. If the short-term immunity hypothesis can be shown to be true, then immunological interference among viruses may be the fundamental driver of the immuno-epidemiology of respiratory disease transmission in the tropics. In temperate countries, where strong wintertime seasonality synchronizes respiratory disease transmission, the interference hypothesis may not be testable due to the short transmission season. In the tropics, where there is no winter to structure the dynamics of respiratory virus transmission, individual viral epidemics may create post-epidemic niches – unfavorable to other respiratory pathogens – by generating temporary waves of nonspecific immunity.

Although a complete forecasting evaluation will require a separate analysis, we can already detect one clear limitation of ILI forecasting methods: that they must be based on future weather predictions which, in our analysis, were bootstrapped from past weather data. Nevertheless, this proved to be a small obstacle in our analysis as, for Ho Chi Minh City, the bootstrapped climate variables yielded accurate predictions of averages for temperature and relative humidity (Figure S5). In other words, it is more likely that higher levels of ILI during a particular period are affected by the average climate behavior during that period, and not by any particular days that have extremes in temperature or relative humidity. This contrasts with the climate mechanisms proposed in temperate zones where it is postulated that the onset of abnormally low absolute humidity is closely associated with the onset of the influenza season^41^. The larger question on climate effects and influenza — why AH, RH, and temperature appear to have different transmission effects in temperate and tropical regions^11,43,59^ — remains to be answered. Much work remains to be done before respiratory disease outbreaks in the tropics can be forecast accurately; our hope is that the non-annual signal identified in this study will help in this endeavor.

A second limitation in the current study design is the lack of age information. We experimented with several different reporting methods (email, log books) for this study, but only the log-book method was able to capture age information consistently. Unfortunately, this method was adopted by a minority of the clinics in our study, and it was not compatible with real-time reporting. The age distribution of ILI cases represents a critical data gap in our study and in other mHealth studies that aim at real-time reporting, as the age distribution could tell us whether the major disease burden skews towards childhood respiratory diseases or general respiratory diseases like influenza. As tropical countries have younger age distributions than temperate countries, this difference may have a profound epidemiological effect on differences in ILI dynamics between temperate and tropical zones, as well as the proportion of ILI cases that are caused by influenza versus other respiratory viruses.

The public health value of our mHealth reporting system is that ILI results can be fed back in real time to participating physicians and the community of health professionals in Ho Chi Minh City. Real-time ILI trends from our study are publicly available and updated daily. The two key questions raised by our study are (*i*) to what extent the transmission of non-influenza respiratory viruses in the tropics is a potential driver of complex multipathogen transmission system, and (*ii*) whether it is useful to attempt the timing of influenza vaccination in an epidemiological scenario where influenza epidemics occur irregularly. We aim to investigate the first of these questions by introducing more respiratory virus diagnostics into our study. The second question can be evaluated with a mathematical model of influenza epidemiology, but will necessitate a longer influenza time series and a better understanding of the key drivers of influenza virus dynamics in tropical settings.

## 4 Materials and Methods

### Influenza-like illness data

In August 2009, a participatory epidemiology study was established in Ho Chi Minh City, Vietnam, in collaboration with the Hospital for Tropical Diseases in Ho Chi Minh City (HCMC) and with permission from the Ho Chi Minh City Department of Health. Participating outpatient clinics report the daily number of total patients seen, the daily number of patients meeting the European CDC definition of influenza-like illness^60^, and the number of hours each clinic was open. To encourage enrollment and reduce dropout, clinics are advised to send daily reports by standard mobile phone short messaging system (SMS) text messages; reporting with log books and email is also available. SMS messages are automatically passed to a text-parsing and data-cleaning system that was set up and is still actively managed by the Oxford University Clinical Research Unit (OUCRU) in HCMC. Every day, ILI reports are manually approved by a qualified project team member at OUCRU; on approval they are automatically entered into a mySQL database that holds all data points for the study. A small number of clinics (about 8%) did not use SMS reporting (by their request) and instead emailed ILI numbers to the project team or wrote them down in a daily logbook provided by OUCRU. As part of the data processing pipeline, reports by email or logbook were regularly merged into the main mySQL database.

Community engagement meetings were run for the first several years of the study to distribute and explain the study protocol, and a basic leaflet outlining the goals of the study and the reporting methodology was distributed to interested physicians. All documents were translated into Vietnamese, and annual reports and ILI trends were fed back to the clinics on a regular basis. A total of 63 clinics were enrolled in the initial study period (August 2009 – December 2015). Clinics that reported frequent zeros (>50%), or withdrew too early (contributed <200 reports) were not considered for the analysis. In May 2012 a new study component was launched for 24 clinics that agreed to periodic collection of naso-pharyngeal (NP) swabs so that a subset of ILI patients could be molecularly confirmed as positive or negative for influenza virus. A swabbing schedule was made at random every year, so that each clinic would be visited an approximately equal number of times; one or two clinics were selected for swabbing each week. The numbers of NP swabs collected each week depended on the numbers of ILI cases presenting at the clinics as well as patient consent. The research protocol was approved by the Oxford Tropical Research Ethics Committee at the University of Oxford and by the Scientific and Ethical Committee of the Hospital for Tropical Diseases in Ho Chi Minh City.

### Molecular Confirmation

Respiratory specimens (nasal/throat swabs) were collected from ILI patients at outpatient clinics, transported the same day on ice to OUCRU, and stored in -80C freezers for a maximum of 3 months before RNA extraction and Influenza A and B PCR testing. All specimens were tested by real-time PCR using primers, probes, and reagents recommended by the World Health Organization (WHO) and the Centers for Disease Control and Prevention (CDC). Sequences of probes and primers used can be referred to in Table S1.

Viral RNA was extracted from 140uL of a patient’s specimen to attain a final elution volume of 50uL. The extraction was carried out using a MagNA Pure 96 automated system (Roche Applied Science) with the MagNA Pure 96 DNA and viral NA Small Volume Kit (Roche; Cat ID. 06543588001), and the MagNa Pure 96 System Fluid (Roche; Cat ID. 05467578001).

Template RNA from the viral extract was used for cDNA synthesis using the LightCycler 480 RNA Master Hydrolysis Probes (Roche; Cat ID. 04991885001). The cDNA products were then amplified in a real-time RT-PCR procedure carri ed out by a LightCycler instrument (Roche Applied Science). Each reaction had a total volume of 20uL containing 5uL of the viral RNA extract, 1X of RNA Master Hydrolysis Probes, 3.25mM of Mn(OAc)_2_, 1X of enhancer solution, 0.2uM of Influenza A/B probes, 0.8uM of Influenza A/B forward primers, 0.8uM of Influenza A/B reverse primers, and water. Equine Arteritis Virus (EAV) was used as an internal control, and included in each reaction with 0.04uM of EAV probes, 0.2uM of EAV forward primers, and 0.2uM of EAV reverse primers. Thermal cycling conditions were set up as follow: reverse transcription at 58C for 20 min, enzyme inactivation at 95C for 5 minutes, and 45 cycles of 95C for 15 seconds, 55C for 30 seconds, and 72C for 20 seconds. Fluorescent signals were measured by LightCyler software, at wavelengths between 465 nm and 510 nm for Influenza A and B.

### Climate Data

Data on daily mean temperature (T) and relative humidity (RH) were collected from Weather Underground for Ho Chi Minh City, Vietnam (http://www.wunderground.com) from the beginning of 2000 till the end of 2015. Absolute humidity (AH) was calculated using relative humidity and temperature:

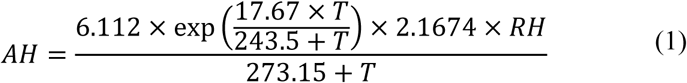

The series of daily climate data were smoothed with a 15-day moving average before being used in our analyses.

### Time series detrending and standardization

A total of 28 regularly reporting clinics (those who reported at least 200 reports from 2010-2015 and reported positive ILI numbers at least half of the time) were included in the time series analysis. A 29th clinic that met these inclusion criteria was removed for quality control reasons. The ILI data of 2009 were not used in the analysis due to the small number of reporting clinics during the first five months of the study. Each clinic’s time series was converted to a z-score scale by computing the z-score of each ILI percentage inside a 12-month moving window (centered at the calculated data point), thus removing long-term trends in the data; we verified that window sizes of 6, 9, 15, and 18 months did not have any qualitative effects on the overall ILI trends. The daily z-scores were averaged across clinics and smoothed using a 15-day window to construct the ILI z-score time series that we used in our subsequent analysis (see Figure S1 for effects of different smoothing windows).

The time series was validated by verifying that is was not white noise (p-value < 10^−15^, Box-Ljung test) and by showing that the majority of individual clinics had a higher correlation to the aggregate time series than would be expected if reporting were random (Figure S2).

### Statistical analysis and forecasting

Periodicity and frequency decomposition in the smoothed ILI trend were assessed with a standard auto-correlation function (ACF) and a Discrete Fourier Transform (DFT). The ILI z-score time series was regressed (linear link function) onto linear and non-linear variants of the climate variables (T, RH, AH, √T, √RH, √AH, T^2^, RH^2^, and AH^2^) to determine which non-linear effects were present, as there is some evidence of non-linear effects of climate on ILI^61^. In addition, a time-dependent fixed effect *α*_j_ mimicking the dominant periodicity identified by the ACF (here, 206 days) was included on the right-hand side of the regression equation. Twenty-one *α_j_* were allowed for in the model, meaning that periodicity in the system is modeled with a piecewise constant function taking 21 different values during a full period of 206 days. This is equivalent to having 21 fixed-effect terms in a regression, each multiplied by an indicator variable describing whether that data point belongs to that period, ensuring that only one fixed-effect term is added at a time. The piecewise constant function has an advantage over the sinusoidal approach traditionally used in epidemiological analyses because the stepwise nature of the *α_j_* allows the periodicity in the system to take any shape determined by the data and does not require that the forcing function to be sinusoidal or continuous.

The non-annual cycle, T, √RH, and RH were the explanatory terms according to the Akaike Information Criterion (AIC) using the stepwise regression approach in R (*step*() function). The ILI z-scores were then regressed onto the non-annual cycle, T, √RH, and RH, and lagged versions of these climate variables, extending back five weeks in the past. The same stepwise regression approach (*step*() function in R) using the AIC was used to remove regression terms that did not add explanatory power. The selected regression equation is

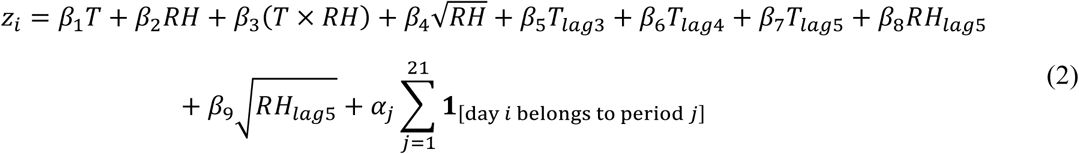

To determine if the regression approach offers any predictability in the system, we inferred the regression coefficients and the time-dependent fixed effects using the first three years of data from January 1st 2010 to December 31st 2012, and we compared the predicted and real ILI trends for 2013–2015. The *median prediction error* was defined simply as the median of all of the absolute differences between the predicted z-score time series and the real z-score time series. We varied the size of the training set to determine how many years of data would be needed to achieve robustness in predictability (Figure S4).

### Bootstrapping climate data

To test the robustness of this prediction to changes in the annual climate cycle and the system’s intrinsic (dominant) cycle identified by the ACF (206 days), we removed the annual trend in the climate cycle with a smoothing-by-bootstrapping approach and we artificially varied the length *c* of the intrinsic non-annual cycle. To create a bootstrap-smoothed climate time series, we defined the climate variables for each time point at *t_bss_* in 2010-2015 as a random sample taken during 2000-2015 and within *d* calendar days of *t_bss_* (see Figure S5). As *d* increases, the annual structure of the climate cycle gradually vanishes. Two hundred bootstrapped time series were created (for each climate variable), for each cycle length *c*, and for each climate subsampling window *d*. For each (*c*, *d*) pair, regression (onto each of the 200 bootstrapped time series separately) and prediction (using each bootstrapped series of 2013-2015 separately) were re-performed, and the median prediction error was plotted to determine if changing assumptions about the length of the intrinsic cycle or the strength/amplitude of the climate data had a detrimental effect on predictability in our system. Mean prediction errors are shown in Figure S7.

All sampling, bootstrapping and statistical analyses were done in R (version 3.2.1, Vienna, Austria).

## Acknowledgements

We are very grateful to all the participating clinicians in this study, to the study nurses Nguyen Thi Kim Cuong, Tran Thi Thao Ly, Tran Thi Anh Tuyet, and Do Ngoc Dung who managed swab collection, and to Professor Tran Tinh Hien who helped introduce us to the first clinicians participating in this study. HML, TDN, NTDN, DNV, NNQM, JEB, GET, DTHT, and HV are funded by Wellcome Trust grant 089276/B/09/7. MFB, NTH, NHTV, TTNT, are NTLT are funded by the Wellcome Trust 098511/Z/12/Z. ST is funded by Wellcome Trust Clinical Fellow (097465/B/11/Z). AW is funded by a James S. McDonnell Foundation postdoctoral fellowship. AW and COB are funded by Models of Infectious Disease Agent Study program (cooperative agreement 1U54GM088558) and acknowledge support from the Wellcome Trust Sustaining Health Grant (106866/Z/15/Z).

## Author Contributions

MFB and JF designed the study. HML and AW analyzed the data. HML designed and implemented the real-time surveillance website www.ili.vn. NTH and NTLT recruited participating clinicians and managed study operations and communication. TDN designed the database and text parsing tools. NTDN was responsible for management and quality control of the data. ST and DNV helped in data analysis and interpretation of results. NTH, NHTV, and TTNT performed molecular diagnostics. PTT, NNQM, JEB, COB, TVN, NVVC, GET, DTHT, and HV helped interpret the data, especially variability in reporting trends. PTT, DTHT, and HV led study reevaluation at various points to ensure high enrollment and improve data quality. MFB, HML, and AW wrote the first draft of the paper.

